# Evaluation of Loop-Mediated Isothermal Amplification (LAMP) for Rapid Detection of *Campylobacter jejuni*

**DOI:** 10.1101/2023.08.03.551860

**Authors:** Junior Caro-Castro, Fiorella Orellana-Peralta, Diana Flores-Leon, Evans Cuch-Meza, Ronnie G. Gavilán, Willi Quino

## Abstract

In recent years, the worldwide incidence of diarrheal diseases caused by *Campylobacter jejuni* has been increasing, causing large-scale outbreaks in developing countries, which added to the capacity of the microorganism to cause Guillain-Barré syndrome (GBS), increasing the need for rapid and timely diagnosis to stop and prevent outbreaks. In this study, a loop-mediated isothermal amplification (LAMP) assay based on the *cdtC* gene was developed and evaluated for the rapid detection of *C. jejuni*, using 91 strains for the standardization and validation. The LAMP assay was compared to whole-genome sequencing as the gold standard. The LAMP assay for *C. jejuni* showed a sensitivity of 100% (CI: 90.94 - 100%), a specificity of 100% (CI: 89.56 - 100%), a positive predictive value of 100% (CI: 90.94 - 100%) and a negative predictive value of 100% (CI: 89.56 - 100%). The assay demonstrated strong agreement between the LAMP assay and genomic sequencing (kappa value = 1). The LAMP assay based on the *cdtC* gene is a method that provides reliable and rapid results, with high sensitivity and specificity for the identification of *C. jejuni*, and is considered a suitable alternative for the diagnosis of diarrheal infections by this pathogen in low-income countries.

## INTRODUCTION

*Campylobacter* species are the leading cause of human foodborne gastroenteritis, whose incidence has increased worldwide, reporting 19.5 and 64.9 cases per 100,000 population in the USA **[1]** and the European Union **[2]**, respectively. In addition, the prevalence of *Campylobacter* in low- and middle-income countries is usually high, although in many of them, the exact infection rates are not known due to deficiencies in the epidemiological study of this pathogen **[3]**.

*Campylobacteriosis* is usually sporadic and self-limited; however, antimicrobial therapy is recommended when patients are infants, immunocompromised, or have other comorbidities. The infectious dose of *Campylobacte*r is usually very low **[4]**, being capable of colonizing the gastrointestinal tract of humans, birds, and wild and domestic mammals, including food-producing animals, as well as having been recovered from contaminated soil, surface water, and groundwater **[5]**. Although the genus *Campylobacter* is composed of 32 species, the majority of infections are associated with *C. jejuni* **[6]**.

*C. jejuni* not only causes gastrointestinal problems, but is also capable of causing an autoimmune condition that affects the nervous system due to cross-reactions between the antigens produced during an infection by this bacterium, called Guillain-Barré syndrome (GBS). This disease is characterized by peripheral nerve disorder, leading to progressive muscle weakness, altered reflexes, and sensory abnormalities **[7]**. GBS cases are considered a major public health problem due to short- and long-term patient complications, since 25% of patients with this disease generally require mechanical ventilation, and their prognosis depends on the type of GBS and prompt patient care **[8]**. This scenario increases the need for timely diagnosis of these infections to avoid complications in affected patients.

An example of the complex public health repercussions of *C. jejuni* infections occurred in Peru in recent years. Reports of the incidence of *Campylobacter* causing gastroenteritis were limited and underestimated for a long time, especially due to the difficulty in isolating and characterizing this pathogen in low-resource areas **[9]**, indicating a 13.3% prevalence in children under the age of 12 years **[10]**, 16.7% of positive carcasses and 26.7% of positive blinds from animals for human consumption **[11]**. However, the abrupt increase in GBS cases between 2018 and 2020 forced the declaration of an epidemiological emergency situation and national health priority, whose efforts made it possible to identify the pathogen causing this disease, a clonal strain of *C. jejuni* ST-2993 that was found in several regions of the country. This scenario demonstrated the lack of epidemiological surveillance of *C. jejuni* in developing countries **[12]**.

Currently, there are several investigations on the genetic bases of *C. jejuni* associated with its virulence as a diarrheal pathogen **[13]** and related to GBS **[14]**, which are used to design molecular diagnostic methodologies for *Campylobacter* species **[15]**. Likewise, the development of methodologies that can be transferred to rural or low-income settings for the accurate detection of pathogens such as *C. jejuni* is proposed, which would allow obtaining more accurate data on the prevalence of diarrheal pathogens in low-resource countries **[16]**. In this context, the design and validation of a molecular method based on loop-mediated isothermal amplification (LAMP) for the detection of *C. jejuni* is proposed, which can be used by reference laboratories and primary care health centers. Therefore, the objective of this study is the development and evaluation of a LAMP assay to detect *C. jejuni* associated with diarrheal diseases and Guillain-Barré syndrome.

## METHODS

### LAMP primer design

Primers for LAMP detection of *C. jejuni* were designed to align with the *C. jejuni* reference genome NCTC 11168 (GBA: NC_002163.1). Primers were designed using LAMP design software (Premier Biosoft International, USA) and were synthesized by Macrogen, Inc. (Korea), whose set consisted of: FIP (Forward Inner Primer), BIP (Backward Inner Primer), F3 (Forward Outer Primer), B3 (Backward Outer Primer), LF (First Forward Loop) and LB (First Backward Loop).

### Samples included in the study

Ninety-one bacterial strains recovered under the genomic surveillance of enteropathogens by the National Reference Laboratory of Clinical Bacteriology, Instituto Nacional de Salud (INS) of Peru were selected **(see Supplementary information 1 online)**. Twenty-eight strains belonged to *C. jejuni* ST-2993, 21 to *C. jejuni* from other sequence types, 21 to *C. coli*, 20 to *Salmonella* Infantis and one to *Escherichia coli*.

### Bacterial culture

*Campylobacter* strains cryopreserved at -80°C were cultured on blood base agar (BD BBL™, USA) with 5% sheep blood, under microaerophilic conditions (85% nitrogen, 10% carbon dioxide and 5% oxygen) at 42 °C for 48 hours [17]. Non-*Campylobacter* strains were cultured on trypticase soy agar (Becton Dickinson, France) and incubated at 37 °C for 24 hours.

### DNA extraction

Bacterial DNA extraction was performed using the PureLink Genomic DNA Mini Kit extraction kit (Invitrogen, USA), following the manufacturer’s recommendations. The concentration and quality of the extracted DNA was determined using a DS-11 FX spectrophotometer/fluorometer (DeNovix, USA).

### LAMP Optimization

LAMP assays were developed with the Warm Start Colorimetric LAMP 2X Master Mix (DNA & RNA) (New England Biolabs, USA), using a mixture of LAMP oligonucleotides at a final concentration of 1.6 μM FIP and BIP; 0.2 μM of oligonucleotides F3 and B3; and 0.4 μM of LF and LB. A reaction mix with a total volume of 25 μL was prepared, containing 12.5 μL of Warm Start Colorimetric LAMP 2X Master Mix, 2.5 μL of LAMP oligonucleotide mix, 5 μL of RNase-free molecular grade water, and 5 μL of DNA. The reaction temperature was 65 °C, using three incubation times: 35, 40 and 45 min in a Thermocycler (Applied Biosystem, USA). LAMP reaction products were evaluated by electrophoresis. Reactions were considered positive if the LAMP products turned from a wine-red to yellow color, as well as a ladder band pattern on agarose gel after electrophoresis.

### Limit of detection of the LAMP assay

Analytical sensitivity was carried out using DNA obtained from a *C. jejuni* strain, with an initial concentration equivalent to 10.7×10^6^ DNA copies in 5 μL. The sample was logarithmically diluted in the range from 10^−1^ to 10^−8^ and evaluated in triplicate. The limit of detection (LoD) was determined by identifying the lowest concentration detected in the assay equivalent to the DNA copy number.

### Analytical specificity of the LAMP assay

The specificity of the LAMP assay was carried out using the DNA of the 91 clinical strains included in this study, all at a concentration of 10.7×10^6^ DNA copies in 5 μL.

### Statistical analysis

The sensitivity and specificity of the LAMP assay were estimated using whole-genome sequencing (WGS) as the reference standard with a 95% confidence interval (95% CI). Concordance of the results obtained by WGS and each of the LAMP assays based on Cohen’s kappa statistic was obtained using the Stata v15.0 statistical program (Stata Corporation, USA). The sensitivity, specificity and linearity plots were constructed with GraphPad Prism v7 (GraphPad Inc., USA).

## RESULTS

### Evaluation of LAMP primers

The primers designed in this study were evaluated with the NCBI Primer-BLAST tool. The sequences of the four sets of designed primers **(Table 1)** were compared with the sequences of the genomes of *C. jejuni, C. coli* and *S*. Infantis described in the NCBI **(see Supplementary information 1 online)**. Based on the *in silico* analysis, primers from set 3 obtained 100% identity with the *cdtC* gene inside the *C. jejuni* reference genome NCTC 11168 and other *C. jejuni* genomes recovered from NCBI such as *C. jejuni* ST-2993 6.897-2019 (GBA: JAGJUO000000000) and *C. jejuni* 1.519-2016 (GBA: JAGTNP000000000), not aligned with negative controls such as *C. coli* 1.776-2017 (GBA: JAGTKF000000000) and *S*. Infantis 1.010-2014 (GBA: GCA_012939885) **(Fig. 1)**.

**Table 1.**
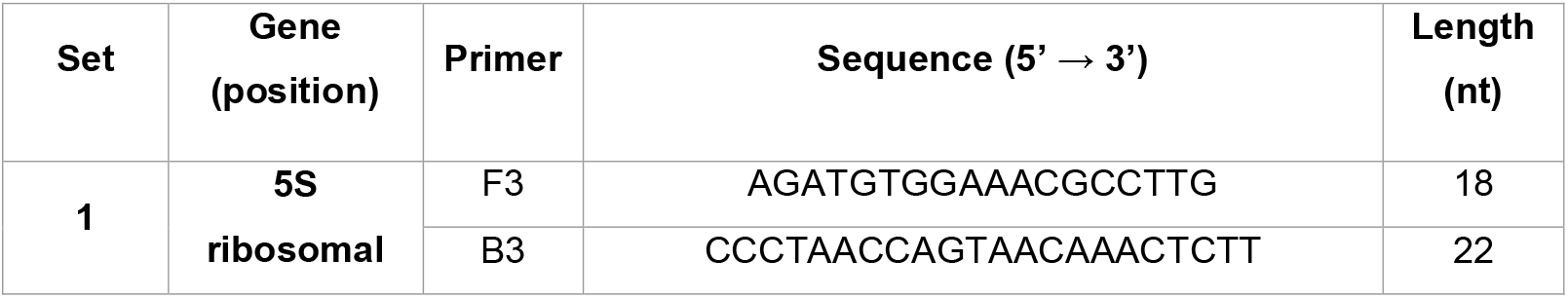
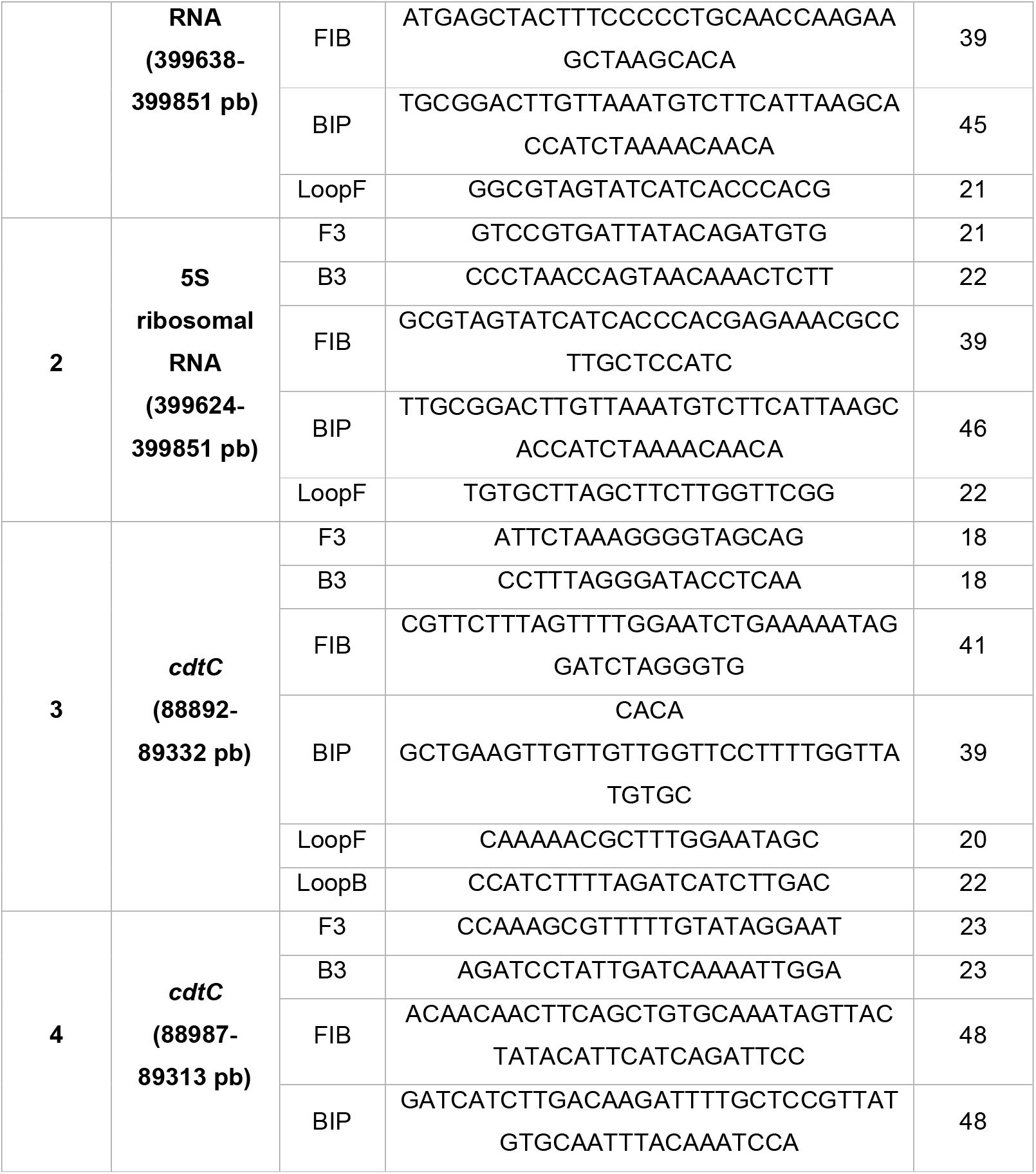
LAMP primers designed for detection of *C. jejuni*.

**Fig. 1.**
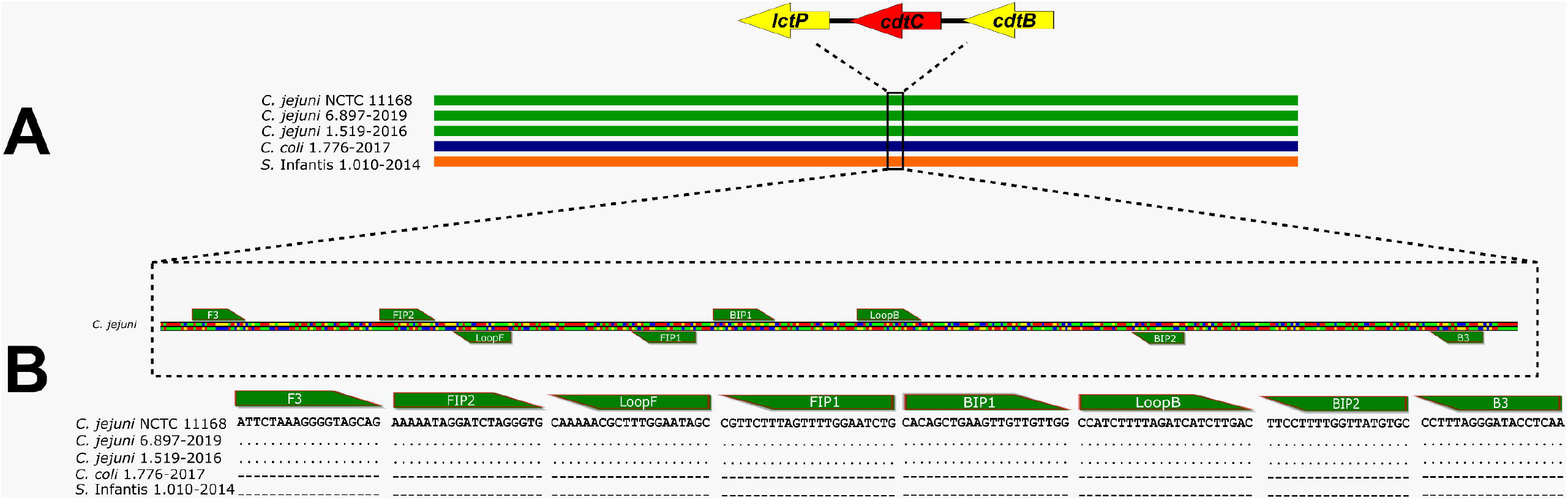
*In silico* analysis of set 3 primer alignment for the *C. jejuni* LAMP assay. **A**. Genetic map of the complete genome of *C. jejuni, C. coli*, and *S*. Infantis **B**. Alignment of primers from set 3 designed for LAMP with *C. jejuni, C. coli* and *S*. Infantis.

*In silico* analysis was corroborated by the *in vitro* LAMP assays, obtaining primers from sets 1 and 2 that amplified both *C. jejuni* and *C. coli*, while primers from set 4 did not amplify with any of the samples evaluated **(Fig. 2)**. Therefore, primers from set 3 were selected to continue with the optimization and validation of the LAMP assay.

**Fig. 2.**
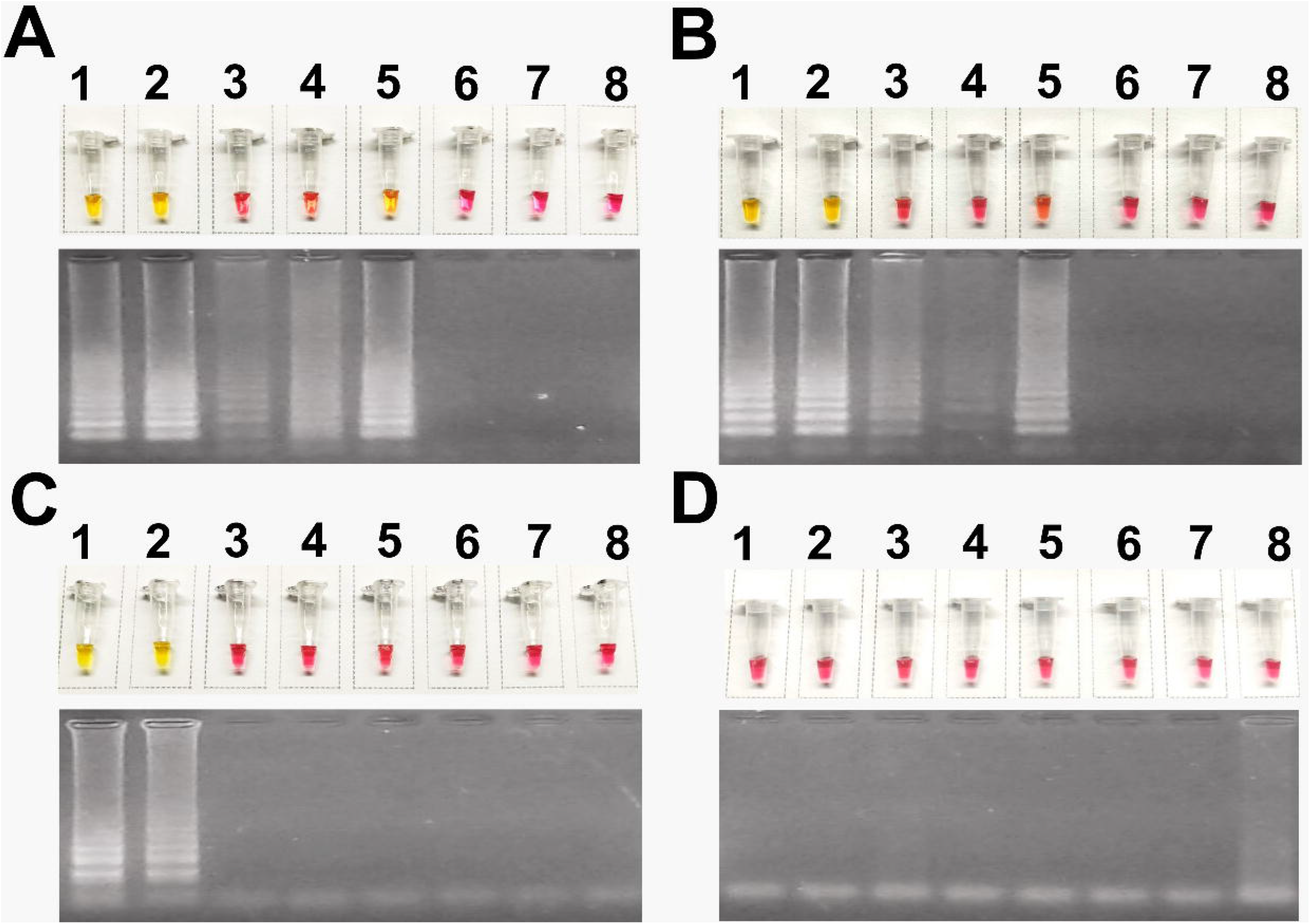
LAMP assay using the four sets of primers designed. **A)** Set 1, **B)** Set 2, **C)** Set 3, **D)** Set 4. The LAMP reaction is visualized at the top of each section, while the agarose gel amplification product is visualized at the bottom. **1)** *C. jejuni* ST-2993, **2)** *C. jejuni* of another genotype, **3-5)** *C. coli*, **6)** *S*. Infantis, **7)** Negative control, **8)** Not template control.

### LAMP Optimization

In the three incubation times evaluated, a positive result was observed for the *C. jejuni* samples of all genotypes evaluated, while *C. coli* and *S*. Infantis strains gave a negative result, which was corroborated by the change in the colorimetric reagent as well as by the observation of amplification products; however, at 35 and 40 minutes, some of the positive samples were not well defined in terms of the change from wine-red to yellow; therefore, the incubation temperature at 45 minutes was considered the best for the LAMP assay. **(Fig. 3)**.

**Fig. 3.**
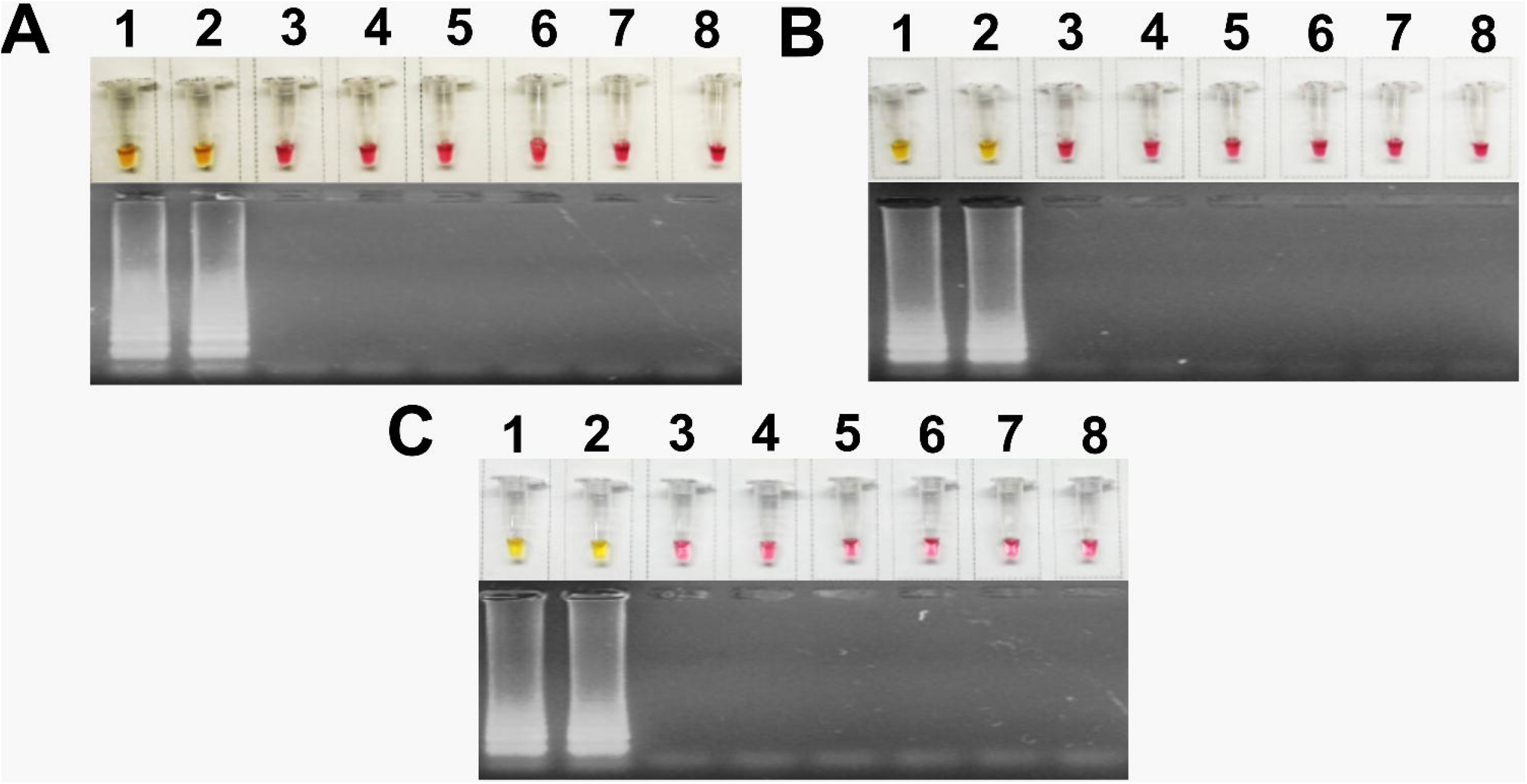
LAMP optimization for different reaction times **A)** 35 minutes, **B)** 40 minutes, **C)** 45 minutes. The LAMP reaction is visualized at the top of each section, while the agarose gel amplification product is visualized at the bottom. **1)** *C. jejuni* ST-2993, **2)** *C. jejuni* of another genotype, **3-5)** *C. coli*, **6)** *S*. Infantis, **7)** Negative control, **8)** Not template control.

### Limit of detection of *C. jejuni* by LAMP

The LAMP technique was able to detect *C. jejuni* samples up to 10^−2^ dilution, indicating that the limit of detection (LoD) of the technique is up to a concentration of 107 000 DNA copies, equivalent to 0.0188 ng or 1.88 pg/μl, allowing the detection of low concentrations of genetic material. The results were confirmed by the presence of DNA bands up to 10^−2^ dilution in the agarose gel **(Fig. 4)**.

**Fig. 4.**
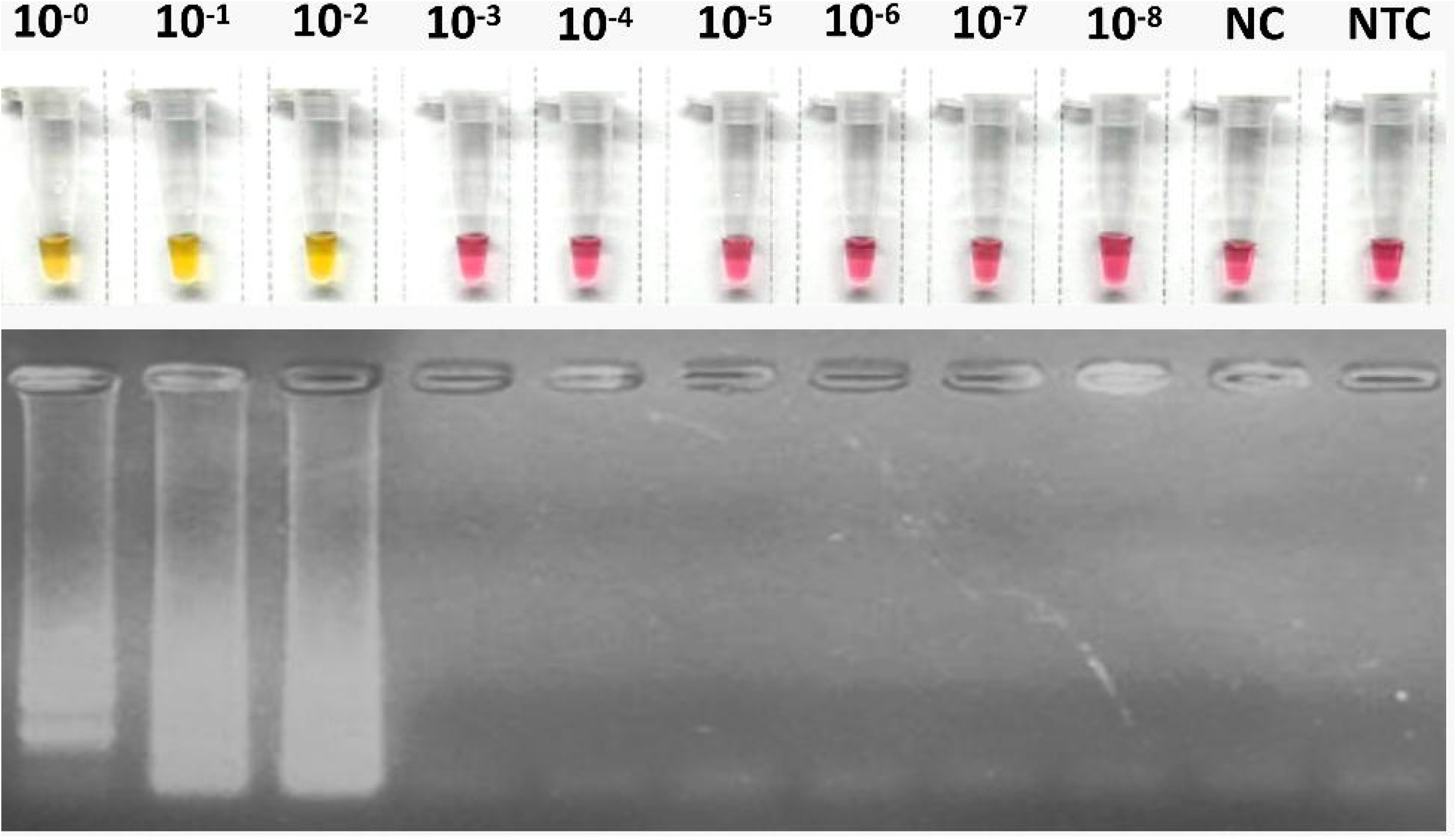
Limit of detection (LoD) of sample 1 for the detection of the *cdtC* gene of *C. jejuni*. The LAMP reaction is visualized at the top, while the agarose gel amplification product is visualized at the bottom.

### LAMP Level of Agreement for *C. jejuni*

The LAMP results of the *cdtC* gene were compared with the results of WGS, detecting a total of 49 true positives and 42 true negatives in a cohort of 91 samples **(Fig. 5)**. In the comparative analysis of WGS with the LAMP assay, no false negatives were observed, reaching a specificity of 100% (90.94% - 100%) and a sensitivity of 100% (89.56% - 100%), with a positive predictive value of 100% (90.94% - 100%) and a negative predictive value of 100% (89.56% - 100%). From the concordance analysis of the results of WGS compared with the LAMP assay for the *cdtC* gene, there is a high concordance of 1 between WGS and LAMP according to Cohen’s kappa statistic.

**Fig. 5.**
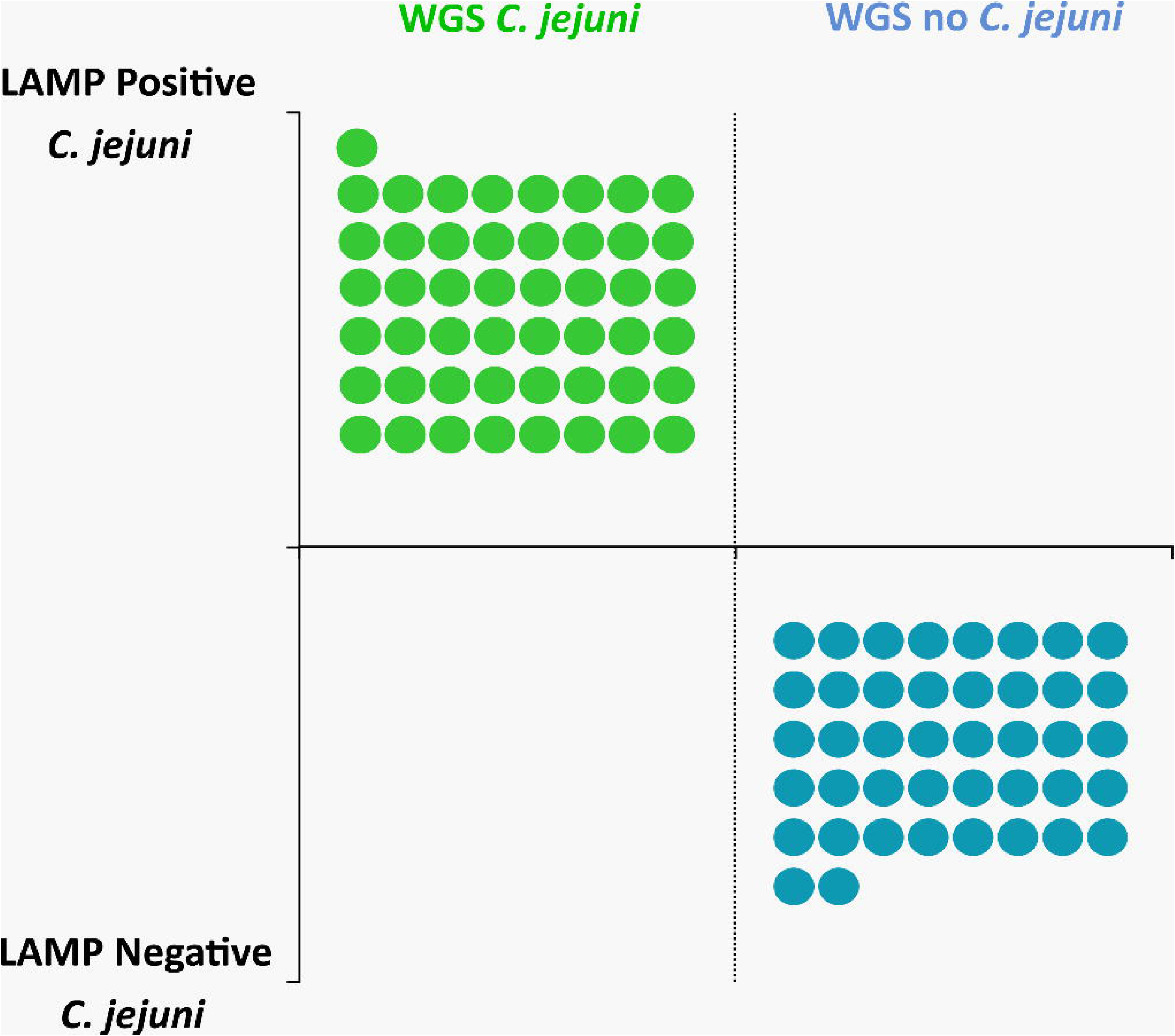
Linearity chart comparing LAMP positive/negative samples and their detection based on *C. jejuni* species-based whole-genome sequencing.

## DISCUSSION

*C. jejuni* is one of the main etiological agents causing watery diarrhea in Peru, mainly affecting children, which adds to the microbial resistance reported by some studies **[11]** and the ability of some of its genotypes to cause nervous disorders such as SGB **[18]**, making it a microorganism of priority importance in public health **[19]**. Culture and biotyping have long been used for clinical reporting of *C. jejuni*; however, due to the metabolic properties of these bacteria, its culture is fastidious, especially due to the dependence on nutritional supplements and the long incubation period **[20]**. For these reasons, the use of molecular techniques has been proposed, which do not depend on a high concentration of microorganisms and allow a faster diagnosis than traditional techniques **[21]**; however, due to the high cost of equipment and reagents, their implementation in low-income areas is difficult. For this reason, the development of low-cost and portable detection protocols such as LAMP for *C. jejuni* has been proposed **[15]**.

Based on BLAST, set 3 of designed primers allowed the specific amplification of a region corresponding to the *cdtC* gene inside the *C. jejuni* genome. In this regard, several markers used for the detection of the *C. jejuni* species by LAMP are found in the literature, such as the *gufA* gene **[22]**, the oxidoreductase gene **[23]** and the *hipO* gene **[16, 24]**, in addition to the *cdtC* gene **[25]** found here. Most of these genes synthesize essential molecules for cell maintenance, which is why their expression is constitutive in any strain of *C. jejuni*, with the exception of the *cdtC* gene, which is involved in the synthesis of cytolethal distending toxin, a factor of characteristic virulence of pathogenic *C. jejuni* **[13]**. On the other hand, no references to LAMP assays aimed at detecting genotypes in *C. jejun*i or other *Campylobacter* species were found, which would indicate the complexity at the time of searching for exclusive targets in each of them.

The set of primers used allowed us to obtain results by establishing the best conditions for amplification at 65 °C for 45 minutes. The best-established reaction temperature coincides with that specified by the brand of reagent used for the LAMP assay; however, the reaction time is slightly longer than specified. In this regard, it is known that most commercially used pH indicators, such as phenol red, are highly sensitive to the presence of protons produced in the LAMP reaction, which are released as byproducts of the incorporation of a deoxynucleoside triphosphate into the nascent DNA by polymerase action, which allows a color change easily detectable by the human eye **[26]**. However, the pH can also be affected by the initial concentration of DNA and the presence of low concentration inhibitors, which slow down the pH change of the indicator using short incubation times, often observing an incomplete change from wine-red to orange, affecting the interpretation of a LAMP result **[27]**. For this reason, 45 minutes was established as the best incubation time because at this temperature, the change from wine-red to yellow is clearly observed, regardless of the initial concentration of the sample or the presence of inhibitors that are not usually measured in routine practice.

Regarding the limit of detection (LoD), the LAMP assay allowed the detection of a sample that has 107000 copies of DNA, equivalent to 1.88 pg/μl, a low concentration of genetic material that can be easily recovered by conventional extraction such as boiling or using phenol-chloroform method **[28]**, and comparable to the reported by Quyen *et al*. **[23]**, who were also able to detect *C. jejuni* in samples with a concentration of up to 1 pg/μl of DNA, but using another marker, the oxidoreductase gene. Although studies with LoD values that detect less than 1 pg/μl of DNA can be found, most of them depend on the use of a fluorometer to perform very low measurements **[24, 25]**, which, as previously emphasized, affects the implementation of this technique in low-income areas such as most of the regions of South America, Africa and Asia. On the other hand, Babu *et al*. **[15]** standardized a colorimetric LAMP protocol that allowed detection of *Campylobacter* up to a concentration of 10 fg/μl of DNA; however, this protocol was only validated for detection at the genus level, but not at the species level.

Finally, 100% sensitivity and 100% specificity were obtained when running the LAMP for *C. jejuni*, values higher than those obtained with other markers such as the oxidoreductase gene, with 100% sensitivity and 97.9% specificity **[23]**, or the *gufA* gene, with 98.5% sensitivity and 97.4% specificity **[22]**. It is necessary to highlight that most of these studies were carried out for the detection of *C. jejuni* in DNA recovered from food samples, which is why their values may be affected. On the other hand, our sensitivity and specificity values are comparable to those achieved in other studies with the same marker, such as that carried out by Kreitlow *et al*. **[25]** who also reached 100% in both parameters, however, the method developed by these researchers depends on a fluorometer to measure the color change during the assay, and although this equipment is portable, it still increases the costs of implementation in low-income areas.

A major limitation in the LAMP assay of this study is the need to use high-quality DNA obtained from pure culture. Strains of the *Campylobacter* genus are usually sensitive to various environmental parameters, with oxygen being the main factor to take into account for the recovery of colonies in enriched culture media, since it has been observed that the efficiency of recovery of microorganisms in samples transported and stored for several days and exposed to oxygen is low or null **[29]**. The use of molecular techniques helps to overcome this limitation, allowing the detection of *Campylobacter* in a nonviable or viable but nonculturable state **[30]**; however, the LAMP method proposed here has only been standardized and validated with DNA obtained from pure culture. Therefore, additional tests are required to verify the efficiency of detection of *C. jejuni* in DNA obtained from the total sample. Some studies have validated the detection of C. *jejuni* from total DNA from stools of infected patients; however, several false negatives were reported compared to results from standard PCR **[16]**. On the other hand, in the studies that used LAMP assays to detect *C. jejuni* in food samples, when comparing their results with those obtained when testing with DNA from pure culture, it was observed that the sensitivity and specificity of the assay were negatively affected **[22, 25]**.

In conclusion, the LAMP assay for *C. jejuni* optimized and validated in this study will allow the molecular detection of the presence of this pathogen in a fast and timely way, and, thanks to its easy handling, low cost and high effectiveness, can be implemented in low-income countries, thus strengthening the epidemiological and molecular surveillance of *C. jejuni* to avoid future outbreaks.

## Abbreviations

BLAST: Basic Local Alignment Search Tool
CI: Confidence interval
DNA: Deoxyribonucleic acid
GBA: GenBank Access
GBS: Guillain-Barré syndrome
INS: Instituto Nacional de Salud
LAMP: Loop-mediated isothermal amplification
LoD: limit of detection
NC: Negative control
NCTC: National Collection of Type Cultures
NTC: Not template control
ST: Sequence Type
WGS: Whole-genome sequencing.

## Acknowledgements

We thank the members of the National Reference Laboratory of Clinical Bacteriology/INS for their technical assistance. This work was supported by the project “Evaluation of Loop-mediated isothermal amplification (LAMP) for the rapid detection of *Campylobacter jejuni* ST2993 associated with Guillain Barré syndrome in Peru during the period 2022-2023”, approved by Institutional Review Boards of Instituto Nacional de Salud of Peru according to D.R. N°261-2022-OGITT/INS.

## Author contributions

JCC, FOP and ECM led on the data analysis and drafted the manuscript. JCC and FOP conducted molecular analyses. JCC, FOP and WQ were involved in the interpretation of the data. DFL, RGG and WQ critically revised the manuscript. All authors read and approved the final manuscript.

## Data availability

All data generated or analyzed during this study are included in this published article. The datasets generated and/or analyzed during the current study are available in the NCBI/Genbank repository, whose accession numbers are indicated in Supplementary information 1 online.

## Ethics declarations

This study was conducted within the framework of the research project “Evaluation of Loop-mediated isothermal amplification (LAMP) for the rapid detection of *Campylobacter jejuni* ST2993 associated with Guillain Barré syndrome in Peru during the period 2022-2023” which was approved by the Instituto Nacional de Salud of Peru (D.R. N°261-2022-OGITT/INS). All procedures and methods were performed in accordance with ethical standards of the Declaration of Helsinki or comparable relevant guidelines and regulations. The approval of an informed consent was waived due to the retrospective nature of this study by the Institutional Committee of Research and Ethics (IRB) of the Instituto Nacional de Salud of Peru, in accordance with the national legislation and the institutional requirements for Public Health Surveillance.

