## Additional file 1 for "Evaluation of Loop-Mediated Isothermal Amplification (LAMP) for Rapid Detection of *Campylobacter jejuni*"

**Table S1.** Genomes used in the design of LAMP primers for *C. jejuni*

| N° | Strain name | Species | ST | Year | GenBank Access number |
|---|---|---|---|---|---|
| 1 | 1.470-2010 | *C. jejuni* | 52 | 2010 | JAGTKV000000000 |
| 2 | 1.514-2010 | *C. jejuni* | 6247 | 2010 | JAGTMK000000000 |
| 3 | 1.516-2010 | *C. jejuni* | 607 | 2010 | JAGTLC000000000 |
| 4 | 1.584-2010 | *C. jejuni* | 6091 | 2010 | JAGTMG000000000 |
| 5 | 1.588-2010 | *C. jejuni* | 8310 | 2010 | JAGTMX000000000 |
| 6 | 1.048-2011 | *C. jejuni* | 5789 | 2011 | JAGTLS000000000 |
| 7 | 1.088-2011 | *C. jejuni* | 4722 | 2011 | JAGTLQ000000000 |
| 8 | 1.109-2011 | *C. jejuni* | 6091 | 2011 | JAGTMB000000000 |
| 9 | 1.110-2011 | *C. jejuni* | 5789 | 2011 | JAGTLT000000000 |
| 10 | 1.134-2011 | *C. jejuni* | 6091 | 2011 | JAGTMD000000000 |
| 11 | 1.137-2011 | *C. jejuni* | 10717 | 2011 | JAGTND000000000 |
| 12 | 1.144-2011 | *C. jejuni* | 862 | 2011 | JAGTLE000000000 |
| 13 | 1.150-2011 | *C. jejuni* | 607 | 2011 | JAGTLA000000000 |
| 14 | 1.151-2011 | *C. jejuni* | 5758 | 2011 | JAGTLR000000000 |
| 15 | 1.152-2011 | *C. jejuni* | 6177 | 2011 | JAGTMI000000000 |
| 16 | 1.160-2011 | *C. jejuni* | 5789 | 2011 | JAGTLU000000000 |
| 17 | 1.161-2011 | *C. jejuni* | 8117 | 2011 | JAGTML000000000 |
| 18 | 1.209-2011 | *C. jejuni* | 8310 | 2011 | JAGTMQ000000000 |
| 19 | 1.263-2011 | *C. jejuni* | 8117 | 2011 | JAGTMM000000000 |
| 20 | 1.265-2011 | *C. jejuni* | 8310 | 2011 | JAGTMR000000000 |
| 21 | 1.266-2011 | *C. jejuni* | 8310 | 2011 | JAGTMT000000000 |
| 22 | 1.554-2011 | *C. jejuni* | 8310 | 2011 | JAGTMV000000000 |
| 23 | 1.562-2011 | *C. jejuni* | 407 | 2011 | JAGTKY000000000 |
| 24 | 1.581-2011 | *C. jejuni* | 8310 | 2011 | JAGTMW000000000 |
| 25 | 1.582-2011 | *C. jejuni* | 5789 | 2011 | JAGTLY000000000 |
| 26 | 1.066-2012 | *C. jejuni* | 10577 | 2012 | JAGTMZ000000000 |
| 27 | 1.134-2012 | *C. jejuni* | 8310 | 2012 | JAGTMN000000000 |
| 28 | 1.141-2012 | *C. jejuni* | 8310 | 2012 | JAGTNW000000000 |
| 29 | 1.235-2012 | *C. jejuni* | 3720 | 2012 | JAGTLP000000000 |
| 30 | 1.278-2012 | *C. jejuni* | N/A | 2012 | JAGTNG000000000 |
| 31 | 1.546-2012 | *C. jejuni* | N/A | 2012 | JAGTNZ000000000 |
| 32 | 1.618-2012 | *C. jejuni* | 6091 | 2012 | JAGTMH000000000 |
| 33 | 1.637-2012 | *C. jejuni* | 52 | 2012 | JAGTKW000000000 |

| | | | | | |
|---|---|---|---|---|---|
| 34 | 1.761-2012 | *C. jejuni* | 7356 | 2012 | JAGTNT000000000 |
| 35 | 1.054-2013 | *C. jejuni* | 1036 | 2013 | JAGTLI000000000 |
| 36 | 1.072-2013 | *C. jejuni* | 6091 | 2013 | JAGTMA000000000 |
| 37 | 1.198-2013 | *C. jejuni* | 6091 | 2013 | JAGTME000000000 |
| 38 | 1.208-2013 | *C. jejuni* | 5789 | 2013 | JAGTLV000000000 |
| 39 | 1.354-2013 | *C. jejuni* | 5789 | 2013 | JAGTLW000000000 |
| 40 | 1.022-2014 | *C. jejuni* | 607 | 2014 | JAGTKZ000000000 |
| 41 | 1.086-2014 | *C. jejuni* | N/A | 2014 | JAGTNA000000000 |
| 42 | 1.087-2014 | *C. jejuni* | 862 | 2014 | JAGTLD000000000 |
| 43 | 1.159-2014 | *C. jejuni* | 8310 | 2014 | JAGTMO000000000 |
| 44 | 1.617-2014 | *C. jejuni* | 2114 | 2014 | JAGTLM000000000 |
| 45 | 1.677-2014 | *C. jejuni* | 1036 | 2014 | JAGTKD000000000 |
| 46 | 1.979-2014 | *C. jejuni* | 137 | 2014 | JAGTKX000000000 |
| 47 | 1.017-2015 | *C. jejuni* | 1233 | 2015 | JAGTKU000000000 |
| 48 | 1.018-2015 | *C. jejuni* | 1233 | 2015 | JAGTLL000000000 |
| 49 | 1.163-2015 | *C. jejuni* | 8310 | 2015 | JAGTMP000000000 |
| 50 | 1.183-2015 | *C. jejuni* | 6177 | 2015 | JAGTMJ000000000 |
| 51 | 1.265-2015 | *C. jejuni* | 8310 | 2015 | JAGTMS000000000 |
| 52 | 1.356-2015 | *C. jejuni* | 10618 | 2015 | JAGTNI000000000 |
| 53 | 1.357-2015 | *C. jejuni* | 8310 | 2015 | JAGTMU000000000 |
| 54 | 1.040-2016 | *C. jejuni* | 5742 | 2016 | JAGTMY000000000 |
| 55 | 1.043-2016 | *C. jejuni* | 6091 | 2016 | JAGTLZ000000000 |
| 56 | 1.113-2016 | *C. jejuni* | 6091 | 2016 | JAGTMC000000000 |
| 57 | 1.519-2016 | *C. jejuni* | 862 | 2016 | JAGTNP000000000 |
| 58 | 1.520-2016 | *C. jejuni* | 5789 | 2016 | JAGTLX000000000 |
| 59 | 1.522-2016 | *C. jejuni* | 10618 | 2016 | JAGTNP000000000 |
| 60 | 1.489-2016 | *C. jejuni* | N/A | 2016 | JAGTNM000000000 |
| 61 | 1.143-2017 | *C. jejuni* | 3515 | 2017 | JAGTLN000000000 |
| 62 | 1.299-2017 | *C. jejuni* | 9354 | 2017 | JAGTNH000000000 |
| 63 | 1.300-2017 | *C. jejuni* | 6091 | 2017 | JAGTMF000000000 |
| 64 | 1.636-2017 | *C. jejuni* | 10618 | 2017 | JAGTNS000000000 |
| 65 | 1.766-2017 | *C. jejuni* | 1915 | 2017 | JAGTNU000000000 |
| 66 | 1.799-2017 | *C. jejuni* | 862 | 2017 | JAGTLH000000000 |
| 67 | 1.198-2018 | *C. jejuni* | 10246 | 2018 | JAGTNF000000000 |
| 68 | 1.418-2018 | *C. jejuni* | 10604 | 2018 | JAGTNJ000000000 |

| | | | | | |
|---|---|---|---|---|---|
| 69 | 1.496-2019 | *C. jejuni* | 10618 | 2019 | JAGTNN000000000 |
| 70 | 1.506-2018 | *C. jejuni* | 862 | 2018 | JAGTLF000000000 |
| 71 | 1.508-2018 | *C. jejuni* | 607 | 2018 | JAGTLB000000000 |
| 72 | 1.510-2018 | *C. jejuni* | 1036 | 2018 | JAGTLJ000000000 |
| 73 | 1.603-2019 | *C. jejuni* | 8923 | 2019 | JAGTNQ000000000 |
| 74 | 1.577-2018 | *C. jejuni* | 3572 | 2018 | JAGTLO000000000 |
| 75 | 6.897-2019 | *C. jejuni* | 2993 | 2019 | JAGJUO000000000 |
| 76 | 6.1083-2019 | *C. jejuni* | 2993 | 2019 | JAGJUX000000000 |
| 77 | 6.1195-2019 | *C. jejuni* | 2993 | 2019 | JAGJUW000000000 |
| 78 | 6.1196-2019 | *C. jejuni* | 2993 | 2019 | JAGJUV000000000 |
| 79 | 6.1197-2019 | *C. jejuni* | 2993 | 2019 | JAGJUU000000000 |
| 80 | 6.1198-2019 | *C. jejuni* | 2993 | 2019 | JAGJUT000000000 |
| 81 | 1.1279-2019 | *C. jejuni* | 2993 | 2019 | JAGJVB000000000 |
| 82 | 1.1280-2019 | *C. jejuni* | 2993 | 2019 | JAGJVA000000000 |
| 83 | 1.1281-2019 | *C. jejuni* | 2993 | 2019 | JAGJUZ000000000 |
| 84 | 1.1282-2019 | *C. jejuni* | 2993 | 2019 | JAGJUY000000000 |
| 85 | 6.2107-2019 | *C. jejuni* | 2993 | 2019 | JAGJUS000000000 |
| 86 | 6.2108-2019 | *C. jejuni* | 2993 | 2019 | JAGJUR000000000 |
| 87 | 6.2116-2019 | *C. jejuni* | 2993 | 2019 | JAGJUQ000000000 |
| 88 | 6.2122-2019 | *C. jejuni* | 2993 | 2019 | JAGKSA000000000 |
| 89 | 6.2139-2019 | *C. jejuni* | 2993 | 2019 | JAGJUP000000000 |
| 90 | 6.059-2020 | *C. jejuni* | 2993 | 2020 | JAGJUN000000000 |
| 91 | 6.060-2020 | *C. jejuni* | 2993 | 2020 | JAGJUM000000000 |
| 92 | 6.066-2020 | *C. jejuni* | 2993 | 2020 | JAGJUL000000000 |
| 93 | 4.166-2020 | *C. jejuni* | 2993 | 2020 | JAGJUH000000000 |
| 94 | 4.167-2020 | *C. jejuni* | 2993 | 2020 | JAGJUG000000000 |
| 95 | 4.168-2020 | *C. jejuni* | 2993 | 2020 | JAGJUF000000000 |
| 96 | 4.169-2020 | *C. jejuni* | 2993 | 2020 | JAGJUK000000000 |
| 97 | 4.170-2020 | *C. jejuni* | 2993 | 2020 | JAGJUJ000000000 |
| 98 | 4.171-2020 | *C. jejuni* | 2993 | 2020 | JAGJUI000000000 |
| 99 | OBT12377 | *C. jejuni* | 2993 | 2019 | CP059157.1 |
| 100 | OBT12390 | *C. jejuni* | 2993 | 2019 | CP059160.1 |
| 101 | OBT12393 | *C. jejuni* | 2993 | 2019 | CP059159.1 |
| 102 | OBT12386 | *C. jejuni* | 2993 | 2019 | CP059158.1 |

| | | | | | |
|---|---|---|---|---|---|
| 103 | ICDCCJ07001 | *C. jejuni* | 2993 | 2007 | GCA_000184085.1 |
| 104 | ICDCCJ07002 | *C. jejuni* | 2993 | 2007 | GCA_000355825.1 |
| 105 | ICDCCJ07004 | *C. jejuni* | 2993 | 2007 | GCA_000355845.1 |
| 106 | NCTC12851 | *C. jejuni* | 45 | 1990/93 | GCA_900638285.1 |
| 107 | HF5-5-1 | *C. jejuni* | 45 | 2012 | GCA_001951255.1 |
| 108 | CJ677CC086 | *C. jejuni* | 677 | 1999 | GCA_001507225.1 |
| 109 | NCTC13257 | *C. jejuni* | 45 | 1999 | GCA_900638225.1 |
| 110 | FDAARGOS_266 | *C. jejuni* | 583 | n/a | GCA_002209065.1 |
| 111 | CJ677CC012 | *C. jejuni* | 794 | 2007 | GCA_001507265.1 |
| 112 | CJ677CC034 | *C. jejuni* | 794 | 2002 | GCA_001507205.1 |
| 113 | THJ097 | *C. jejuni* | 8071 | 2019 | GCA_024349525.1 |
| 114 | FDAARGOS_1546 | *C. jejuni* | 267 | n/a | GCA_020736145.1 |
| 115 | HF5-7-1 | *C. jejuni* | 45 | 2012 | GCA_001951275.1 |
| 116 | CJ677CC095 | *C. jejuni* | 677 | 2007 | GCA_001507245.1 |
| 117 | 1.252-2015 | *C. coli* | 1055 | 2015 | JAGTKJ000000000 |
| 118 | 1.254-2015 | *C. coli* | 1055 | 2015 | JAGTKK000000000 |
| 119 | 1.260-2015 | *C. coli* | 5123 | 2015 | JAGTKL000000000 |
| 120 | 1.266-2015 | *C. coli* | N/A | 2015 | JAGTKM000000000 |
| 121 | 1.268-2015 | *C. coli* | 830 | 2015 | JAGTKN000000000 |
| 122 | 1.352-2015 | *C. coli* | 902 | 2015 | JAGTKO000000000 |
| 123 | 1.707-2017 | *C. coli* | N/A | 2017 | JAGTKE000000000 |
| 124 | 1.776-2017 | *C. coli* | 825 | 2017 | JAGTKF000000000 |
| 125 | 1.807-2017 | *C. coli* | 825 | 2017 | JAGTKG000000000 |
| 126 | 1.507-2018 | *C. coli* | 8317 | 2018 | JAGTKA000000000 |
| 127 | 1.491-2019 | *C. coli* | 8939 | 2019 | JAGTKS000000000 |
| 128 | 1.497-2019 | *C. coli* | 8939 | 2019 | JAGTJZ000000000 |
| 129 | 1.467-2010 | *C. coli* | N/A | 2010 | JAGTKR000000000 |
| 130 | 1.404-2011 | *C. coli* | N/A | 2011 | JAGTKP000000000 |
| 131 | 1.413-2011 | *C. coli* | 8317 | 2011 | JAGTKQ000000000 |
| 132 | 1.497-2011 | *C. coli* | 8317 | 2011 | JAGTKT000000000 |
| 133 | 1.530-2012 | *C. coli* | N/A | 2012 | JAGTKB000000000 |
| 134 | 1.567-2012 | *C. coli* | 860 | 2012 | JAGTKC000000000 |
| 135 | 1.195-2013 | *C. coli* | N/A | 2013 | JAGTKH000000000 |
| 136 | 1.197-2013 | *C. coli* | N/A | 2013 | JAGTKI000000000 |
| 137 | 1.667-2014 | *C. coli* | 1055 | 2014 | JAGTLK000000000 |

| | | | | | |
|---|---|---|---|---|---|
| 138 | FDAARGOS_1464 | *C. coli* | 45 | n/a | GCA_020149725.1 |
| 139 | 1.197-2015 | *S. Infantis* | 32 | 2015 | GCA_012272245.1 |
| 140 | 1.010-2014 | *S. Infantis* | 32 | 2014 | GCA_012939885.1 |
| 141 | 1.042-2014 | *S. Infantis* | 32 | 2014 | GCA_012939845.1 |
| 142 | 1.068-2014 | *S. Infantis* | 32 | 2014 | GCA_012939865.1 |
| 143 | 1.072-2014 | *S. Infantis* | 32 | 2014 | GCA_012939875.1 |
| 144 | 1.279-2014 | *S. Infantis* | 32 | 2014 | GCA_012939855.1 |
| 145 | 1.346-2014 | *S. Infantis* | 32 | 2014 | GCA_012939945.1 |
| 146 | 1.485-2014 | *S. Infantis* | 32 | 2014 | GCA_012939985.1 |
| 147 | 1.598-2014 | *S. Infantis* | 32 | 2014 | GCA_012272105.1 |
| 148 | 1.206-2015 | *S. Infantis* | 32 | 2015 | GCA_012272245.1 |
| 149 | 1.607-2014 | *S. Infantis* | 32 | 2014 | GCA_012272165.1 |
| 150 | 1.618-2014 | *S. Infantis* | 32 | 2014 | GCA_012272115.1 |
| 151 | 1.645-2014 | *S. Infantis* | 32 | 2014 | GCA_012271965.1 |
| 152 | 1.669-2014 | *S. Infantis* | 32 | 2014 | GCA_012272135.1 |
| 153 | 1.973-2014 | *S. Infantis* | 32 | 2014 | GCA_012272095.1 |
| 154 | 1.990-2014 | *S. Infantis* | 32 | 2014 | GCA_012271975.1 |
| 155 | 1.004-2015 | *S. Infantis* | 32 | 2014 | GCA_012271935.1 |
| 156 | 1.006-2015 | *S. Infantis* | 32 | 2015 | GCA_012272225.1 |
| 157 | 1.011-2015 | *S. Infantis* | 32 | 2015 | GCA_012272195.1 |

**N/A:** Non available
